# Threat-Induced Anxiety During Goal Pursuit Disrupts Amygdala-Prefrontal Cortex Connectivity in Posttraumatic Stress Disorder

**DOI:** 10.1101/614446

**Authors:** Delin Sun, Andrea L. Gold, Chelsea A. Swanson, Courtney C. Haswell, Vanessa M. Brown, Daniel Stjepanovic, VA Mid-Atlantic MIRECC Workgroup, Kevin S. LaBar, Rajendra A. Morey

## Abstract

To investigate how unpredictable threat during goal pursuit impacts fronto-limbic activity and functional connectivity in posttraumatic stress disorder (PTSD), we compared military veterans with PTSD (n=25) versus trauma-exposed Control (n=25). Participants underwent functional magnetic resonance imaging (fMRI) while engaged in a computerized chase-and-capture game task that involved optimizing monetary rewards obtained from capturing virtual prey while simultaneously avoiding capture by virtual predators. The game was played under two alternating contexts – one involving exposure to unpredictable, task-irrelevant threat by randomly occurring electrical shocks, and a nonthreat control condition. Activation in and functional connectivity between the amygdala and ventromedial prefrontal cortex (vmPFC) was tested across threat and nonthreat task contexts with generalized psychophysiological interaction (gPPI) analyses. PTSD patients reported higher anxiety than controls across contexts. Better task performance represented by successfully avoiding capture by predators under threat than nonthreat contexts was associated with stronger left amygdala-vmPFC functional connectivity in controls and greater vmPFC activation in PTSD patients. PTSD symptom severity was negatively correlated with vmPFC activation in trauma-exposed controls and with right amygdala-vmPFC functional connectivity across all participants in the threat relative to nonthreat contexts. The findings showed that veterans with PTSD have disrupted amygdala-vmPFC functional connectivity and greater localized vmPFC processing under threat-modulation of goal-directed behavior, specifically related to successful task performance while avoiding loss of monetary rewards. In contrast, trauma survivors without PTSD rely on stronger threat-modulated left amygdala-vmPFC functional connectivity during goal-directed behavior, which may represent a resilience-related functional adaptation.

## Introduction

Imminent threat elicits fear and is accompanied by phasic fight or flight responses, whereas unpredictable threat is associated with anxiety and sustained hypervigilance and apprehension^1^. Posttraumatic stress disorder (PTSD) is characterized by symptoms of hyperarousal and hypervigilance that produce considerable distress and functional impairment^2^. The most widely adopted behavioral models for studying PTSD are based on fear conditioning and extinction^3,4^, but few studies have examined the brain response associated with the anxiety elicited by the uncertainty resulting from unpredictable threat^5^. Rodents consistently prefer predictable shocks and their associated contexts, and predictability attenuates the negative effects of stress^5^. Rodents exposed to unpredictable threats display a behavioral syndrome akin to PTSD, such as hypervigilance, insomnia, and impaired attention^6,7^. The functional impairment in humans produced by uncertainty is a core feature of anxiety^8^ that is related to PTSD symptoms of avoidance, numbing, and hyper-arousal^9,10^. Intolerance to uncertainty is known to predict subsequent PTSD symptoms following campus shootings^11^. Patients with PTSD who are treated with cognitive-behavioral therapy show decreased startle magnitude to unpredictable threat, and this decline is correlated with a decline in PTSD symptoms^12^. Unpredictable threat, frequently experienced by many veterans during deployment, has been linked to impaired goal-directed processing^13,14^, a component of executive functioning, which is compromised upon returning to the demands of civilian life^15^. A central claim of the *generalized unsafety theory of stress* (GUTS) is that veterans are unable to switch-off or inhibit the default stress response, which becomes dependent on the perception of generalized unsafety rather than actual threat^16^.

It is widely accepted that effective regulation of the amygdala response by the prefrontal cortex (PFC), particularly ventromedial PFC (vmPFC)^17^, is crucial for successfully maintaining goal-directed behavior^18^. We previously reported that non-clinical volunteers exposed to unpredictable threat during a computer gaming task elicited greater functional connectivity of right amygdala with vmPFC^13^. Furthermore, right amygdala functional connectivity with vmPFC positively correlated with successful goal-directed behavior during unpredictable threat. Reduced functional connectivity between amygdala and vmPFC has also been documented in PTSD patients regardless of specific task requirements^19,20^.

Our aim was to investigate the functional effects on fronto-limbic systems, particularly the amygdala and vmPFC, when patients with PTSD are exposed to unpredictable threat (unexpected shocks) while simultaneously balancing competing task demands. In this paradigm^13^, participants face threat from unpredictable, task-irrelevant shocks in some task blocks while navigating a virtual avatar through a maze to pursue moving prey and evade pursuit by predators. Prey capture by the avatar and predator capture of the avatar were motivated by monetary gains and losses, respectively. This task design requires active engagement and rapid ongoing response of the participant that imposes identical demands for tracking behavior across unpredictable threat and nonthreat contexts. We hypothesized that better performance on the *chase-and-capture game* (indexed by either more prey captures or fewer avatar captures) would be associated with greater amygdala-vmPFC functional connectivity in trauma-exposed controls than in PTSD patients.

## Method

### Participants

Twenty-five participants with PTSD and 25 trauma-exposed controls were recruited from a repository of military service members and veterans who served after September 11, 2001^21^. Psychiatric diagnoses were confirmed using the Clinician-Administered PTSD Scale for DSM-5 (CAPS-5)^2^. Exclusion criteria included major Axis I disorders (other than depressive or anxiety disorders), contraindication to MRI, traumatic brain injury, substance dependence, neurological disorders, and age over 65 years. Symptom assessments included Beck Depression Inventory–II (BDI-II)^22^, Alcohol Use Disorders Identification Test (AUDIT)^23^, Drug Abuse Screening Test (DAST)^24^, Child Trauma Questionnaire (CTQ)^25^, State-Trait Anxiety Inventory (STAI)^26^, Traumatic Life Events Questionnaire (TLEQ)^27^, and combat exposure (CES)^28^. Eleven participants with PTSD and no controls took medications listed in **Supplementary Table 1S**. All participants provided written informed consent to participate in procedures reviewed and approved by the Institutional Review Boards at Duke University and the Durham Veterans Affairs Medical Center. Participants were compensated $25/hour and gained a task bonus of $10-25 based on task performance.

### Experimental Procedure

The chase-and-capture task paradigm (**Fig. 1**) was described by Gold et al.^13^. Under the task-irrelevant threat of electrical shock, participants used a joystick to navigate an avatar within a 2D maze to capture prey, which generated monetary rewards, and to evade capture by a predator (avatar capture), which incurred monetary loss. The predator, under software control, followed the minimum path with the goal of capturing the avatar.

**Figure 1.**
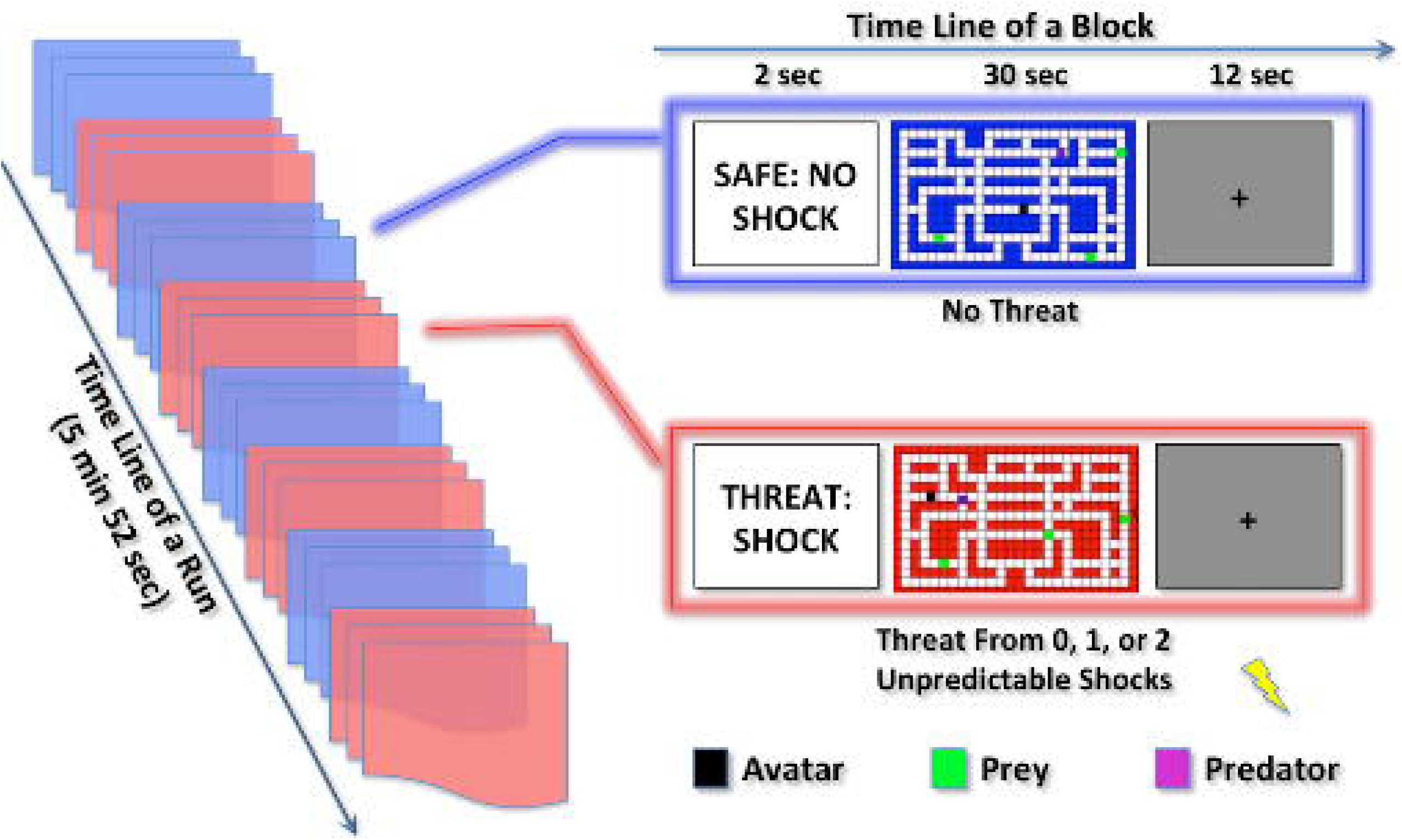
Schematic of the chase-and-capture game paradigm. Participants performed five runs of the task. In each run, there were four threat (represented in red) and four nonthreat (represented in blue) blocks presented in alternating order. In each block, a text cue signaling block type was displayed for 2 seconds, then followed by a computer game lasting for 30 seconds, and ended by a fixation cross exhibiting for 12 seconds. During the computer game, participants were asked to move an avatar (black square) within a 2D maze on the screen through operating joystick to capture prey (green squares) and to avoid capture by a predator (purple square). Prey capture by the avatar and avatar capture by the predator were associated with monetary gain and loss, respectively. No shock was accompanied with the nonthreat block, while there was 0 shock in some threat blocks, and 1 or 2 shocks in the other threat blocks (at least one shock per run, and on average 0.35 shocks per threat block). The onset of a shock was randomized relative to the onset of the embedded block. The participants had no way to distinguish between threat blocks without shock and threat blocks with unpredictable shocks before the shock delivery.

Prior to the task, the intensity of shock was calibrated for each participant according to his/her tolerance threshold (**Supplementary Data**). Participants received detailed instructions and completed a practice run. An adaptively-defined difficulty level was utilized to equilibrate task performance of avatar capture across participants. The task difficulty indexed via predator speed from least difficult (level-1) to most difficult (level-5 where predator speed=avatar speed) was modulated by computer program in the practice run. The median value of difficulty was applied and remained constant throughout the MRI task. Task difficulty did not differ between-groups (PTSD, 4.3±0.9; controls, 4.5±0.7; *t*(48) = −0.719, *p* = 0.475).

In the MRI task, participants performed five runs of four threat blocks and four nonthreat blocks, which were presented in alternating order per run. No shocks were presented during the nonthreat condition, but unconditional shocks would be delivered at random times during the threat condition. Blocks lasted 30 seconds and were separated by a 12-second rest period. A cue at the beginning of each block was presented for 2 seconds to signal an impending threat or nonthreat block. Most threat blocks were allocated 1 or 2 shocks, but occasional blocks had no shocks (≥ 1 shock/run, average=0.35 shocks/block). The onset of a shock was randomized relative to the onset of the embedded block to minimize collinearity in fMRI analyses.

After the experiment, participants self-reported task strategy and anxiety under various contexts: (1) “did you focus more on avoiding predator (=1) or catching prey (=7)?”, (2) “How anxious were you when you entered mazes? 1=not at all to 9=highly anxious”, (3) “how anxious were you when staying in mazes? 1=not at all to 9=highly anxious”, and (4) “how much did you dread being chased by predator? 1=not at all to 9=high dread”.

### Behavioral Data Analyses

Avatar captures and prey captures were recorded every 0.5 second. The average number of prey captures, the average number of avatar captures, and the scores of four self-report questions were each entered into a two-way repeated-measures analysis of variance (ANOVA) with Group (PTSD vs. controls) as a between-subjects factor and Context (threat vs. nonthreat) as a within-subjects factor. Four participants with missing data were excluded from behavioral analyses. Results of post-hoc analyses were corrected by Bonferroni method.

### Imaging Acquisition and Preprocessing

Images were acquired on a 3-Tesla GE scanner equipped with an 8-channel headcoil. T1-weighted whole-brain axial images were obtained with 1-mm isotropic voxels using array spatial sensitivity encoding technique (ASSET) and fast spoiled gradient-recall (3D-FSPGR) (TR/TE/flip angle=8.16-ms/3.18-ms/12°, FOV=256-mm^2^, 1 mm slice thickness, 172 slices, 256×256 matrix). Functional images were obtained using the standard echo-planar pulse sequence (TR/TE/flip angle = 2000-ms/27-ms/60°, FOV=256-mm^2^, 64×64 matrix, 3.8 mm thickness, 34 oblique axial slices, no interslice gap).

Images were preprocessed using the CONN toolbox (https://sites.google.com/view/conn/). After discarding the first three volumes, functional images were slice-time and head-motion corrected, and co-registered to each participant’s structural image. Structural images were segmented, bias corrected and spatially normalized to the Montreal Neurological Institute (MNI) space. The normalization parameters were also applied to normalize the functional images. Finally, functional images were smoothed with an 8-mm FWHM Gaussian kernel.

### Blood Oxygen Level Dependent (BOLD) Activation

General Linear Modelling (GLM) with SPM12 (http://www.fil.ion.ucl.ac.uk/spm/) examined BOLD activation. In the individual-level GLM analyses, the regressors for shock occurrences, threat, nonthreat conditions, and nuisance regressors for six head-motion parameters were modeled and convolved with a canonical hemodynamic response function. The shock regressor was modeled with onset in which the shock was delivered and with duration of 0.6 second. The threat and nonthreat condition regressors were modeled with onset of the cue and with the block duration of 32 seconds (cue=2 seconds, plus maze=30 seconds). High-pass temporal filtering with a cut-off of 128 seconds was employed to remove low-frequency drift.

The group-level statistical inferences were based on mean betas of BOLD responses extracted from the regions of interest (ROI) through MarsBar software (http://marsbar.sourceforge.net). ROIs of bilateral amygdala were defined by the WFU PickAtlas (https://www.nitrc.org/projects/wfu_pickatlas), and the functionally defined vmPFC ROI (peak MNI coordinates 2,40,-12) was from our prior study using this paradigm in non-clinical participants^13^ (**Fig. 2A**). The mean betas of brain activity were entered into a three-way repeated-measures ANOVA with Group (PTSD vs. controls) as a between-subjects factor and Context (threat vs. nonthreat) as well as ROI (left amygdala, right amygdala vs. vmPFC) as within-subjects factors. Post-hoc analyses findings were corrected by Bonferroni method.

**Figure 2.**
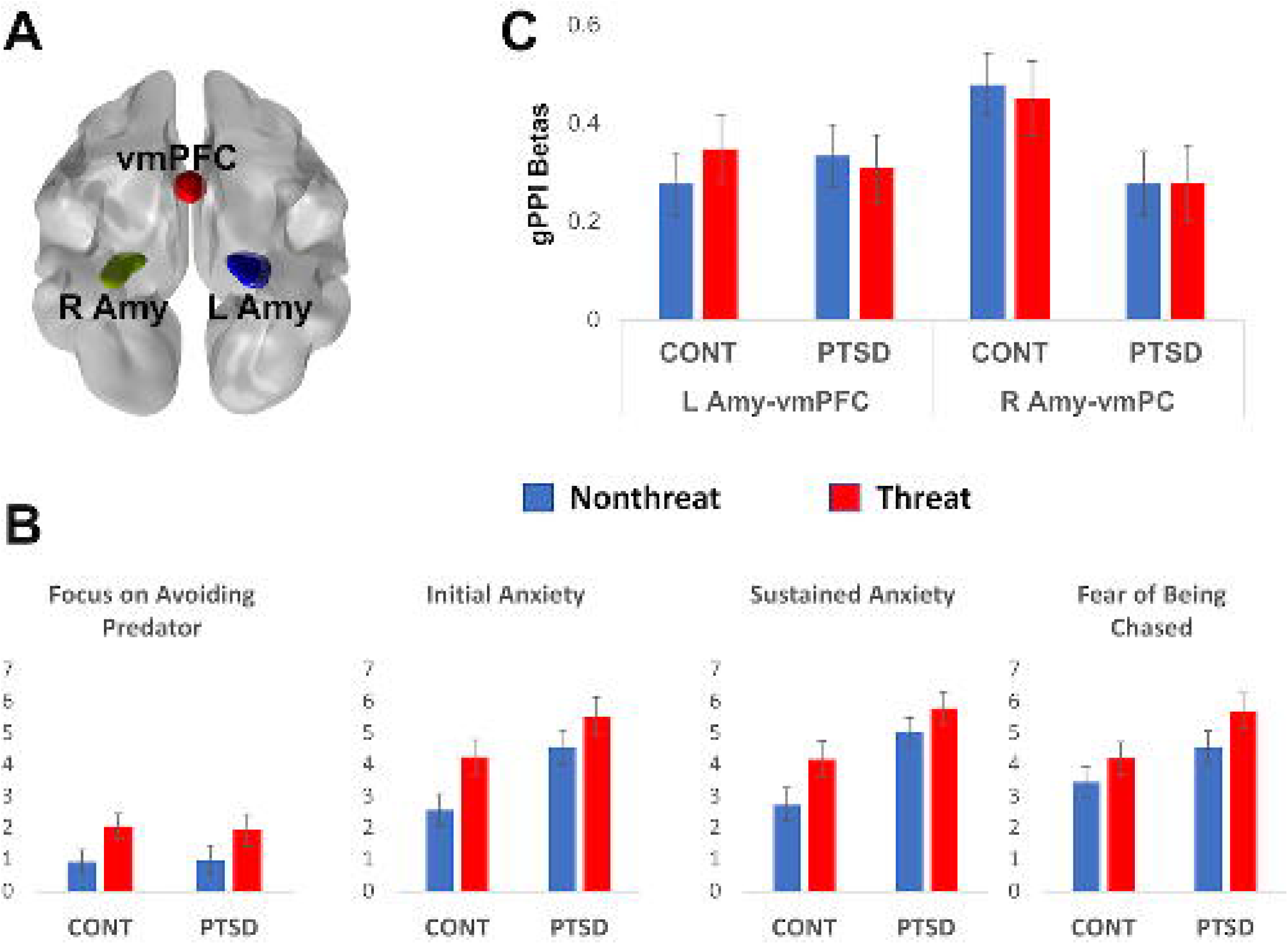
Regions of interest (ROIs), self-report and functional connectivity results. (A) Three ROIs (bottom view): left amygdala (L Amy), vmPFC and right amygdala (R Amy). (B) Self-report results. Threat-vs-nonthreat context elicited enhanced focus on avoiding predator (*F*(1,45) = 7.181, *p* = 0.010), increased initial (*F*(1,44) = 11.221, *p* = 0.002) and sustained anxiety (*F*(1,44) = 7.469, *p* = 0.009), and enhanced fear of being chased by the predator (*F*(1,44) = 7.299, *p* = 0.010). Participants with PTSD compared to controls (CONT) showed greater initial (*F*(1,44) = 6.266, *p* = 0.016) and sustained (*F*(1,44) = 9.937, *p* = 0.003) anxiety, and a trend of significance of enhanced fear to be chased by the predator (*F*(1,44) = 4.035, *p* = 0.051), but non-significant difference in focus on avoiding predator (*F*(1,45) = 0.008, *p* = 0.929). Note: For better understanding, the scores of focus on avoiding predator = 7 - the scores of the first self-report, since the first self-report question was “did you focus more on avoiding predator (=1) or catching prey (=7)?” (C) Controls (*F*(1,24) = 5.821, *p* = 0.048) but not participants with PTSD (*F*(1,24) = 0.988, *p* = 0.660) showed stronger right than left amygdala-vmPFC functional connectivity collapsed across contexts. Error bar denotes standard errors of mean.

The whole brain voxel-wise analyses of threat versus nonthreat and its between-group difference were conducted for exploration purposes only. The results were height-thresholded at *p* < 0.001 and subjected to correction of family wise error (FWE) of *p* < 0.05.

### Generalized Psychophysiological Interaction

Functional connectivity was investigated with the generalized psychophysiological interaction (gPPI) toolbox(https://www.nitrc.org/projects/gppi). We investigated vmPFC correlations with either the left or the right amygdala’s seed time course (physiological regressor) that were significantly modulated by the threat-vs-nonthreat task contrast (an interaction of physiological and psychological regressors). The mean betas of functional connectivity in vmPFC were entered into a three-way repeated-measures ANOVA with Group (PTSD vs. controls) as a between-subjects factor and Seed (left vs. right amygdala) as well as Context (threat vs. nonthreat) as within-subjects factors. Post-hoc analyses findings were corrected by Bonferroni method.

For the BOLD and gPPI analyses, additional ROIs including bilateral inferior frontal gyrus were tested and reported in the **Supplemental material**.

### Brain-Behavior Associations

Pearson’s correlations measured how brain responses (activations or functional connections) to the threat-vs-nonthreat contrast were associated with task performance (average number of prey capture or avatar capture) or clinical measures (PTSD severity indexed by CAPS scores or trauma exposure indexed by TLEQ scores). The brain-clinical associations were conducted in both groups to show the full range of symptom severity, rather than restricting the range to patients. The between-group differences of brain-behavior correlations were compared using the Fisher’s r to z-transformation^29^ and were corrected by Bonferroni method for the number of ROIs (left amygdala, right amygdala vs. vmPFC) or seed-target pairs (left amygdala-vmPFC vs. right amygdala-vmPFC).

We also investigated the correlations between brain responses (activations or functional connections) to either threat or nonthreat context and the corresponding task performance (average number of prey capture or avatar capture).

## Results

### Demographic Information

As shown in **Table 1**, groups did not differ on gender, childhood trauma, substance/alcohol use disorder. PTSD patients compared to controls were slightly older, and had higher levels of trauma exposure, combat exposure, depressive symptoms, and state and trait anxiety. As shown in **Supplementary Data**, after removing the oldest subject in PTSD patients and the youngest subject in controls, the two groups were not significantly different in age, and showed results consistent with the findings reported in the main text.

**Table 1.** Demographic information and clinical measures.

| TEST | CONT | PTSD | PTSD vs. CONT |  |
| --- | --- | --- | --- | --- |
|  | *Mean (SD)* | *Mean (SD)* | *Statistics* | *p-value* |
| **Sex (M/F)** | 22/3 | 22/3 | NS |  |
| **Age** | 38.2(7.8) | 43.7(10.1) | 2.143 | 0.037 |
| **CAPS** | 6.3(6.6) | 46.9(14.7) | 12.633 | <0.001 |
| **BDI-II** | 4.9(6.9) | 18.2(13.2) | 4.438 | <0.001 |
| **CTQ** | 46.0(24.2) | 54.2(24.5) | 1.197 | 0.237 |
| **CES** | 3.5(4.7) | 14.2(12.0) | 4.144 | <0.001 |
| **TLEQ** | 12.7(12.2) | 22.1(10.8) | 2.912 | 0.005 |
| **AUDIT** | 4.0(3.9) | 3.7(2.8) | -0.345 | 0.732 |
| **DAST** | 0.2(0.7) | 0.8(1.2) | 1.817 | 0.077 |
| **STAI-state** | 27.7(8.4) | 40.8(11.5) | 4.578 | <0.001 |
| **STAI-trait** | 29.7(7.9) | 44.5(11.8) | 5.197 | <0.001 |

Note: CONT, trauma-exposed controls; PTSD, PTSD patients; CAPS, Clinician-Administered PTSD Scale reflecting symptoms in the worst 30 day period of subject's life; BDI, Beck Depression Inventory-II; CTQ, Child Trauma exposure questionnaire; CES, Combat Exposure Scale; TLEQ, Traumatic Life Events Questionnaire; AUDIT, Alcohol Use Disorders Identification Test; DAST, Drug Abuse Screening Test; STAI-state, state anxiety of State-Trait Anxiety Inventory; STAI-trait, trait anxiety of State-Trait Anxiety Inventory; NS, no significance.

### Task Performance

The average number of prey capture did not differ by Context (*F*(1,48) = 0.489, *p* = 0.488) or the Group x Context interaction (*F*(1,48) = 0.088, *p* = 0.768), but showed a trend towards significance for Group (*F*(1,48) = 3.164, *p* = 0.082) depicting that controls (mean±SD: nonthreat, 6.229±0.977; threat, 6.271±1.136) captured more preys than participants with PTSD (mean±SD: nonthreat, 5.543±1.507; threat, 5.646±1.674).

The average number of avatar capture did not differ by Group (*F*(1,48) = 0.422, *p* = 0.519) and Context (*F*(1,48) = 1.103, *p* = 0.299), but revealed a Group x Context interaction (*F*(1,48) = 5.823, *p* = 0.020). Post-hoc analyses found no between-context difference in participants with PTSD (mean±SD: nonthreat, 1.403±0.785; threat, 1.303±0.707; *t*(24) = −1.017, *p* = 0.590), but showed a trend towards significance in which controls were less captured in nonthreat than threat context (mean±SD: nonthreat, 1.093±0.638; threat, 1.350±0.893; *t*(24) = 2.246, *p* = 0.068).

### Self-Report Assessments

As shown in **Fig. 2B**, The main effect of Context showed that threat relative to nonthreat context elicited enhanced focus on avoiding predator relative to capturing prey (*F*(1,45) = 7.181, *p* = 0.010), increased initial (*F*(1,44) = 11.221, *p* = 0.002) and sustained anxiety (*F*(1,44) = 7.469, *p* = 0.009), and enhanced fear of being chased by the predator (*F*(1,44) = 7.299, *p* = 0.010). The main effect of Group showed that participants with PTSD compared to controls showed greater initial (*F*(1,44) = 6.266, *p* = 0.016) and sustained (*F*(1,44) = 9.937, *p* = 0.003) anxiety, and a trend of enhanced fear to be chased by the predator (*F*(1,44) = 4.035, *p* = 0.051), but non difference in focus on avoiding predator (*F*(1,45) = 0.008, *p* = 0.929). No Group x Context interactions were observed (*F*-values < 0.8, *p*-values > 0.4).

### Brain Activation

The significant main effect of Context showed that threat-vs-nonthreat context elicited larger activation across ROIs in both groups (*F*(1,48) = 5.330, *p* = 0.025). The significant main effect of ROI (*F*(2,88) = 9.259, *p* = 0.002) indicated larger activation in both left (*F*(1,48) = 10.944, *p* = 0.006) and right (*F*(1,48) = 9.113, *p* = 0.012) amygdala than vmPFC, whereas no significant difference between left and right amygdala (*F*(1,48) = 0.24, *p* > 1).

The main effect of Group (*F*(1,48) = 0.002, *p* = 0.961), Group x Context interaction (*F*(1,48) = 2.141, *p* = 0.150), Group x ROI (*F*(2,96) = 0.191, *p* = 0.719), Group x Context x ROI (*F*(2,96) = 0.134, *p* = 0.827) and Context x ROI (*F*(2,96) = 1.638, *p* = 0.205) were non-significant.

Whole-brain voxelwise analyses showed that threat compared to nonthreat context was associated with lower activation in the bilateral lingual gyrus (peak Z value = 3.95, cluster size = 780 voxels, peak coordinates x=−4, y=−70, z=16). Neither main effect of Group nor the Group x Context interaction was detected significant.

### Amygdala-vmPFC Functional Connectivity

Independent of task context, the significant Seed x Group interaction (*F*(1,48) = 6.515, *p* = 0.014) indicated larger right than left amygdala-vmPFC functional connectivity in controls (*F*(1,24) = 5.821, *p* = 0.048, **Fig. 2C**) but not in participants with PTSD (*F*(1,24) = 0.988, *p* = 0.660).

The main effect of Group (*F*(1,48) = 1.200, *p* = 0.279), Group x Context interaction (*F*(1,48) = 0.318, *p* = 0.576), Seed (*F*(1,48) = 2.044, *p* = 0.159), Context (*F*(1,48) = 0.020, *p* = 0.889), and Seed x Context interaction (*F*(1,48) = 1.054, *p* = 0.310) were all non-significant.

### Brain-Performance Associations

For the threat-vs-nonthreat contrast, as shown in **Fig. 3A**, better performance indexed by fewer avatar captures was accompanied with larger vmPFC activation in PTSD patients (*R* = 0.551, *p* = 0.004) but not in controls (*R* = −0.259, *p* = 0.211), and the two correlations were significantly different (Fisher’s *z* = 2.935, *p* = 0.009). As shown in **Fig. 3B**, better performance (i.e., fewer avatar captures) also related to stronger left amygdala-vmPFC functional connectivity in controls (*R* = 0.436, *p* = 0.029) but not in PTSD patients (*R* = −0.223, *p* = 0.285), and the two correlations were significantly different (Fisher’s *z* = 2.301, *p* = 0.042).

**Figure 3.**
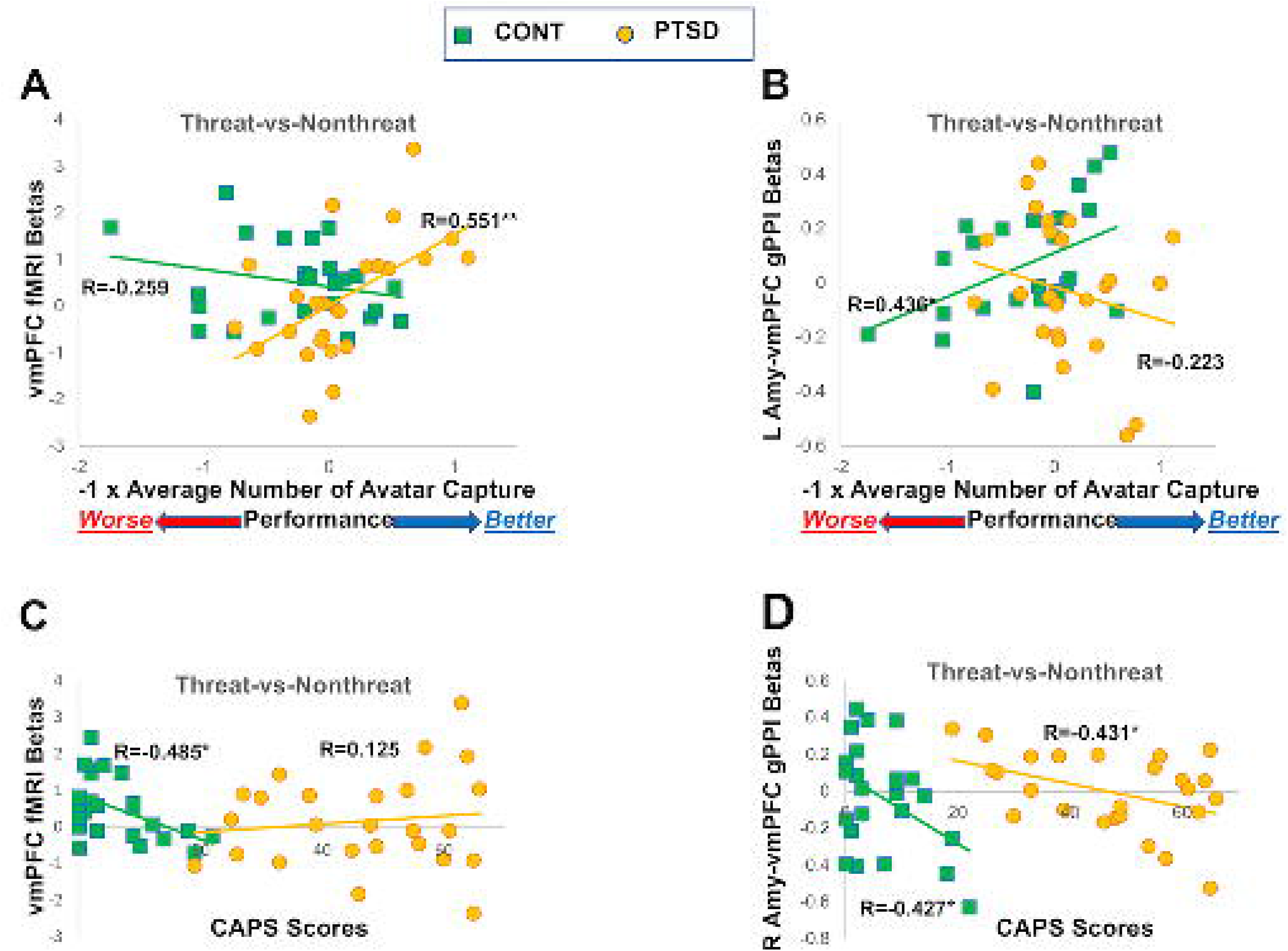
Brain-performance and brain-clinical associations. For the threat-vs-nonthreat contrast, better performance (i.e. fewer avatar captures) is associated with (A) larger vmPFC activation in PTSD (*R* = 0.551, *p* = 0.004) but not in controls (*R* = −0.259, *p* = 0.211; two correlations were significantly different, Fisher’s *z* = 2.935, *p* = 0.009), and (B) stronger left amygdala-vmPFC functional connectivity in trauma-exposed controls (*R* = 0.436, *p* = 0.029) but not in participants with PTSD patients (*R* = −0.223, *p* = 0.285; two correlations were significantly different, Fisher’s *z* = 2.301, *p* = 0.042). For observation purpose only, the x-axis reflects −1 x average number of avatar capture, so that higher values represent better performance. CAPS scores were negatively correlated with (C) vmPFC activity in controls (*R* = −0.485, *p* = 0.014) but not PTSD participant (*R* = 0.125, *p* = 0.553; two correlations were different at a trend level, Fisher’s *z*= −2.173, *p* = 0.090), and (D) right amygdala-vmPFC functional connectivity in both PTSD (*R* = −0.431, *p* = 0.032) and CONT (*R* = −0.427, *p* = 0.033) groups, in response to the threat-vs-nonthreat contrast. *, *p* < 0.05; **, *p* < 0.005.

The number of prey capture did not correlate with either brain activation or functional connections (*p*-values > 0.1). Correlations between performance and brain response in either threat or nonthreat context are reported in **Table 2**.

**Table 2.** Task performance correlates with brain activation per ROI and functional connectivity (gPPI) per seed.

| Performance | Condition | PTSD [R(p)] | CONT [R(p)] | PTSD vs CONT<br>p-value |
| --- | --- | --- | --- | --- |
| ***vmPFC*** |  |  |  |  |
| Prey Capture | Threat | 0.236(0.255) | -0.082(0.695) | 0.283 |
|  | Nonthreat | 0.175(0.404) | -0.304(0.139) | 0.103 |
| Avatar Capture | Threat | -0.261(0.207) | -0.300(0.145) | 0.889 |
|  | Nonthreat | -0.198(0.344) | -0.368(0.070) | 0.537 |
| ***L Amygdala*** |  |  |  |  |
| Prey Capture | Threat | 0.116(0.582) | -0.317(0.122) | 0.140 |
|  | Nonthreat | 0.143(0.495) | -0.514(0.009) | 0.018 |
| Avatar Capture | Threat | 0.157(0.453) | -0.411(0.041) | 0.048 |
|  | Nonthreat | 0.174(0.406) | -0.505(0.010) | 0.015 |
| ***R Amygdala*** |  |  |  |  |
| Prey Capture | Threat | -0.094(0.655) | -0.308(0.134) | 0.457 |
|  | Nonthreat | -0.090(0.668) | -0.398(0.049) | 0.273 |
| Avatar Capture | Threat | -0.074(0.726) | -0.067(0.749) | 0.983 |
|  | Nonthreat | 0.149(0.477) | -0.188(0.368) | 0.259 |
| ***Left Amygdala-vmPFC Functional Connectivity*** |  |  |  |  |
| Prey Capture | Threat | 0.158(0.450) | -0.251(0.226) | 0.168 |
|  | Nonthreat | -0.001(0.996) | -0.053(0.802) | 0.864 |
| Avatar Capture | Threat | -0.011(0.959) | -0.124(0.555) | 0.706 |
|  | Nonthreat | 0.130(0.535) | -0.077(0.714) | 0.490 |
| ***Right Amygdala-vmPFC Functional Connectivity*** |  |  |  |  |
| Prey Capture | Threat | 0.209(0.317) | -0.263(0.203) | 0.110 |
|  | Nonthreat | -0.087(0.680) | -0.055(0.795) | 0.915 |
| Avatar Capture | Threat | 0.172(0.412) | -0.140(0.503) | 0.297 |
|  | Nonthreat | 0.226(0.278) | 0.069(0.744) | 0.626 |

Note: R and p values were based on Pearson correlation within either trauma-exposed controls (CONT) or PTSD patients (PTSD). The between-group comparisons were based on fisher's r to z-transformation, and the *p*-values were uncorrected.

### Brain-Clinical Associations

For the threat-vs-nonthreat contrast, higher CAPS score was accompanied with smaller vmPFC activation in controls (*R* = −0.485, *p* = 0.014, **Fig. 3C**) but not in PTSD patients (*R* = 0.125, *p* = 0.553), and the two correlations showed a trend-level difference (Fisher’s *z*= −2.173, *p* = 0.090). Higher CAPS score was also associated with smaller right amygdala-vmPFC functional connectivity in both controls (*R* = −0.427, *p* = 0.033) and PTSD patients (*R* = −0.431, *p* = 0.032), **Fig. 3D**, whereas the two correlations were not significantly different (Fisher’s *z*= 0.016, *p* = 0.987). CAPS score was related with neither brain activation in amygdala (*p*-values > 0.1) nor the left amygdala-vmPFC functional connectivity in both groups (*p*-values > 0.1).

No brain response correlated with the TLEQ score in either group (*p*-values > 0.1).

For the aforementioned statistical outputs, supplemental statistical models involving covariates for gender, childhood trauma, alcohol abuse, and drug abuse yielded consistent findings (**Supplementary Data**).

## Discussion

Here we investigated how task-irrelevant, unpredictable threats impacted frontolimbic circuitry during goal pursuit in recent war veterans with PTSD and trauma-exposed controls. We used an engaging chase-and-capture game that involved both obtaining monetary rewards and avoiding monetary losses based on task performance. Thus, the task required both vigilance to avoid predators (monetary losses) and pursuit of prey (rewards), which placed demands on attentive, motivational, and cognitive capacities, particularly in the context of task-irrelevant threats. PTSD patients relative to trauma-exposed veterans reported feeling more anxious overall in the task, but both groups reported that the threat context enhanced focus on avoiding predator, increased initial and sustained anxiety, and enhanced fear of being chased by the predator. Consistent with our a priori hypothesis, the results showed that the better performance represented by the successful avoidance of avatar capture by predators was associated with stronger threat-modulated functional connectivity between left amygdala and vmPFC in controls but not in PTSD. Additionally, the successful avoidance of avatar capture by predators was associated with heightened threat-modulated regional activation in the vmPFC in PTSD but not in controls. In addition, we found a negative correlation between PTSD severity and threat-modulated functional connectivity between the right amygdala and vmPFC in both groups, and a negative correlation between PTSD severity and threat-modulated vmPFC activation in controls only.

Threat-modulated association between performance and amygdala-vmPFC functional connectivity differed as a function of PTSD (**Fig 3B**). This is consistent with the dominant theory that PTSD is accompanied with aberrant amygdala-vmPFC circuitry^17^. This study is the first to investigate functional connectivity associated with unpredictable threat induced anxiety in PTSD during goal-directed actions. While the functional connectivity findings in PTSD may reflect impaired top-down inhibition of the amygdala by vmPFC^30^, it may also reflect disrupted bottom-up modulation of vmPFC by the amygdala, or dysregulated bidirectional modulation. In both cases, cortical-subcortical interactions, which have previously been implicated in regulating negative emotions^31^ and maintaining neural functions required for goal-directed behavior^13^, are challenged by unpredictable environmental threats in PTSD. However, gPPI is a correlational method, and advanced, directional modeling methods are needed to uncover the direction of functional connectivity between the amygdala and vmPFC in PTSD. In contrast to the veterans with PTSD, trauma-exposed controls show stronger left amygdala-vmPFC functional connectivity related to predator capture to accomplish the task, which may represent a post-trauma functional adaptation and/or a pre-trauma resilience marker.

The present left-lateralization of threat-modulated task performance association with amygdala-vmPFC functional connectivity in controls is consistent with reports that harm avoidance is associated with left amygdala-vmPFC functional connectivity in healthy subjects^33^. However, risk tolerance^34^ and reward processing^13^ are associated with right amygdala-vmPFC functional connectivity, suggesting a functional dissociation between the two connections. These differences suggest that the amygdala-vmPFC functional connections are lateralized by contextual information, and the downstream implications of threat such as risk, loss, and harm, but also by the potential for profit. The present finding shows that PTSD further modulates this calculus to influence lateralization of amygdala-vmPFC functional connectivity.

Furthermore, PTSD severity was inversely related to threat-modulated functional connectivity between right amygdala and vmPFC in both groups of participants. Optogenetic activation of amygdala afferents to the rodent infralimbic cortex increases anxiety^32^, while inhibiting these afferents has anxiolytic effects. It is possible that veterans with more severe PTSD symptoms in the present study may have inhibited the bottom-up afferents from right amygdala to vmPFC to a larger extent to diminish the interference of anxiety on goal-directed actions. Our finding that brain connectivity during explicit task engagement can predict PTSD symptoms may have potential clinical applications to help in the diagnosis and treatment monitoring of PTSD. This result remains significant when controlling for clinical confounds either by adding the relevant covariates or excluding the affected subjects (**Supplementary Data**).

In contrast to most previously published reports that used fear conditioning paradigms to study phasic anxiety and its neural response, the present study engaged patients with PTSD in a dynamic goal-directed task coupled with exposure to unpredictable threat^35^. In trauma-exposed veterans resilient to PTSD, exposure to unpredictable threat produces sustained subjective anxiety^13^, whereas predictable threat elicits a phasic fear response with a lower level of sustained anxiety^36^. As compared to generalized anxiety disorder, patients with PTSD experience greater sustained anxiety from exposure to unpredictable threat^37^. In fact, the startle reaction to unpredictable threat is a better proxy than predictable threat for treatment success of PTSD and other fear-based disorders such as panic and social anxiety disorder^12^. The persistent inability to perceive safety produces a generalized response following prolonged exposure to unpredictable threat and PTSD^38^. Veterans show greater sensitivity to prediction errors for negative outcomes, and higher PTSD symptom severity is accompanied with lower value-tracking in the amygdala^39^.

Prior research links PTSD to hypoactivation in vmPFC and hyperactivation in amygdala during negative emotion processing^17^. Here we find significant performance-related between-group differences in vmPFC activity but not in amygdala activity. Our participants appear to employ different strategies that impact the utilization of vmPFC, and vmPFC functional connections to amygdala, to effect performance differences. The vmPFC plays a crucial role in fear extinction and extinction retention^17^, and PTSD is associated with decreased vmPFC activation in response to threat or stress^19^. US Special Forces who are resilient to severe trauma display stronger vmPFC activation in response to the expectation of reward ^40^. Furthermore, PTSD patients with robust vmPFC activation are better able to associate previously encountered aversive stimuli with safety than patients with weak vmPFC activation^41^. In line with these previous findings, stronger vmPFC activation might contribute to fulfilling goal-directed tasks by inhibiting conditioned fear responses to unpredictable threats. It is possible that veterans with PTSD place a greater reliance on local processing in the vmPFC to compensate for amygdala-vmPFC functional dysconnectivity.

### Limitations and Strengths

There are two strengths of this study. First, although the PTSD group showed higher levels of trauma exposure, trauma exposure was not related to regional activation or connectivity measures, suggesting the present results cannot be simply explained by magnitude of trauma exposure. Second, the most important findings of the brain-performance and brain-clinical associations are not based on the threat or nonthreat context separately but the threat-vs-nonthreat contrast, which is better to reflect the inner processes of anxiety induced by the unpredictable threat without the confounding effects such as motion. There are also two shortcomings in the present study. First, there is no predictable-threat condition to serve as a control condition. Second, the PTSD patients are slightly older than the trauma-exposed controls, but re-analyses on a subset of participants matched for age, showed comparable results (**Supplementary Data**).

### Conclusion

Veterans with PTSD showed disrupted amygdala-vmPFC functional connectivity and place greater reliance on localized vmPFC processing under threat-modulation of goal-directed behavior, specifically related to successful task performance while avoiding loss of monetary rewards. On the other hand, trauma survivors without PTSD rely on stronger left amygdala-vmPFC functional connectivity during goal-directed behavior that is modulated by threat, which may represent a resilience-related functional adaptation.

## Acknowledgements

This project was supported in part by the Department of Veterans Affairs’ (VA) Mid-Atlantic Mental Illness Research, Education and Clinical Center (MIRECC) of the VA Office of Mental Health Services, the Mid-Atlantic Healthcare Network, and the Office of Research and Development (ORD; 5I01CX000748-01, 5I01CX000120-02). Additional financial support was provided by the National Institute for Neurological Disorders and Stroke (R01NS086885-01A1; Dr. Morey). Work Group Members: Drs. Kimbrel and Dedert were supported by VA Career Development Awards #IK2CX000525 and IK2CX000718, respectively, from the Clinical Science Research and Development (CSR&D) Service. Dr. Van Voorhees was supported by a VA Career Development Award (#5IK2RX001298) from the Rehabilitation Research and Development (RR&D). Dr. Beckham was supported by a VA Research Career Scientist Award (#11S-RCS-009). Dr. Gold was supported by the Intramural Research Program at the National Institute of Mental Health. Research reported in this publication was supported by the Office of the Director, National Institutes of Health under Award Number S10 OD 021480.

