## Supplementary Materials for "Threat-Induced Anxiety During Goal Pursuit Disrupts Amygdala-Prefrontal Cortex Connectivity in Posttraumatic Stress Disorder"

### 1. Shock Calibration

Prior to the chase-and-capture game task, the intensity of shock was calibrated for each participant according to his/her tolerance threshold. Shocks lasting 50 ms were increased in 5-volt increments, beginning from 20 volts, until the participant reported that the shock was “highly annoying, but tolerable.” Shocks were delivered through two electrodes placed on the right ankle below the medial malleolus utilizing the BIOPAC Systems STM200 module with BIOPAC 108A electrode leads (BIOPAC Systems, Goleta, CA). The shock delivery system was grounded through the RF filter panel and was shielded from magnetic interference. Participants were randomly administered 1-4 shocks within a block containing shocks, and they thus could not predict when additional shocks might be delivered.

### 2. BOLD Responses to Shock

Group-level whole brain voxel-wise analyses of shock and the between-group difference of shock was conducted for exploration purposes only. The results were height-thresholded at *p* < 0.001 and subjected to correction of family wise error (FWE) of *p* < 0.05.

Shock elicited stronger activations in several areas including bilateral amygdala, bilateral hippocampus, anterior cingulate cortex, bilateral inferior frontal gyrus/insula. Shock also associated with deactivations in bilateral middle frontal gyrus and bilateral postcentral gyrus.

No significant between-group difference was detected for BOLD responses to shock.

### 3. Behavioral and Brain Results (Model with Covariates)

To investigate whether our findings are confounded by factors including comorbidities and environmental exposures previously reported to be associated with PTSD (9-11), we re-did the analyses reported in the main text by adding gender, childhood trauma exposure (Child Trauma Questionnaire, CTQ) (5), alcohol abuse (Alcohol Use Disorders Identification Test, AUDIT) (3), and drug abuse (Drug Abuse Screening Test, DAST) (4) as covariates. As shown below, re-analyses with co-variates showed comparable results relative to the findings reported in the main text.

#### 3.1 Task Performance

The average number of prey capture was non-significant for Groups (*F*(1,44) = 1.965, *p* = 0.168), Contexts (*F*(1,44) = 0.054, *p* = 0.818), and the Group x Context interaction (*F*(1,44) = 0.001, *p* = 0.997).

The average number of avatar capture was non-significant for Groups (*F*(1,44) = 0.110, *p* = 0.742) and Context (*F*(1,44) = 1.534, *p* = 0.222). The Group x Context interaction was significant (*F*(1,44) = 4.654, *p* = 0.036). However, further analysis showed non-significant between-group differences under either threat (*F*(1,44) = 1.255, *p* = 0.269) or nonthreat (*F*(1,44) = 0.170, *p* = 0.682) context, and non-significant between-context difference in controls (*F*(1,20) = 1.030, *p* = 0.322) and participants with PTSD (*F*(1,20) = 2.213, *p* = 0.152).

#### 3.2 Self-Report Questions

The PTSD participants compared to controls showed increased initial (PTSD: 5.0±0.5, CONT: 3.5±0.5, *F*(1,40) = 4.527, *p* = 0.040) and sustained anxiety (PTSD: 5.4±0.5, CONT: 3.5±0.4, *F*(1,40) = 7.383, *p* = 0.010), as well as a trend of significance of more fear of being chased by the predator (PTSD: 5.1±0.5, CONT: 3.8±0.4, *F*(1,40) = 3.631, *p* = 0.064).

There was also a trend of significance that threat-vs-nonthreat context elicited increased initial anxiety (threat: 4.9±0.4, nonthreat: 3.6±0.4, *F*(1,40) = 2.930, *p* = 0.095)

No other effects were significant (*F*-values < 1, *p*-values > 0.3).

#### 3.3 Brain Activation in ROIs

There was no significant main effect of Group (*F*(1,44) = 0.035, *p* = 0.853) or Context (*F*(1,44) = 0.001, *p* = 0.993). The Group x Context (*F*(1,44) = 2.041, *p* = 0.160), Context x ROI (*F*(2,88) = 0.259, *p* = 0.715), ROI x Group (*F*(2,88) = 0.204, *p* = 0.708), and Context x ROI x Group (*F*(2,88) = 0.042, *p* = 0.925) interactions were all non-significant. The main effect of ROI was significant (*F*(2,88) = 3.738, *p* = 0.049). However, further comparisons between any pair of ROIs did not survive corrections (*p*-values > 0.1).

#### 3.4 Amygdala-vmPFC Functional Connectivity

There was a significant Seed x Group interaction (*F*(1,44) = 6.371, *p* = 0.015). Further analyses showed that, controls versus PTSD participants showed larger right amygdala-vmPFC functional connectivity (*F*(1,44) = 5.419, *p* = 0.049) across contexts, whereas there was no significant between-group difference in the left amygdala-vmPFC functional connectivity (*F*(1,44) = 0.069, *p* = 0.794 uncorrected). All of the other effects were non-significant (*F-value*s < 3.5, *p-value*s > 0.06).

#### 3.5 Brain-Performance Associations

For the threat-vs-nonthreat contrast, better performance (i.e. fewer avatar captures) was accompanied with larger vmPFC activation in PTSD participants (*R* = -0.544, *p* = 0.011) but not in controls (*R* = 0.363, *p* = 0.106), and the two correlations were significantly different (*z*= -3.284, *p* = 0.003). Better performance (i.e. fewer avatar captures) also related to stronger left amygdala-vmPFC functional connectivity in controls (*R* = -0.531, *p* = 0.013) but not in PTSD participants (*R* = 0.146, *p* = 0.527), and the two correlations were significantly different (*z*= -2.450, *p* = 0.028).

There were also trends of significance that the average number of prey capture positively correlated with vmPFC activation (CONT: *R* = 0.432, *p* = 0.051; PTSD: *R* = -0.072, *p* = 0.758) and negatively correlated with left amygdala-vmPFC functional connectivity (*R* = -0.403, *p* = 0.070; PTSD: *R* = 0.114, *p* = 0.622) in controls but not in PTSD participants. The correlations were not significantly different between groups (*p*-values > 0.1).

#### 3.6 Brain-Clinical Associations

For the threat-vs-nonthreat contrast, higher CAPS score was accompanied with smaller vmPFC activation in controls (*R* = -0.438, *p* = 0.047) but not in PTSD participants (*R* = 0.095, *p* = 0.683), whereas the two correlations were not significantly different (*p*-value > 0.1). Higher CAPS score was also associated with smaller right amygdala-vmPFC functional connectivity in PTSD participants (*R* = -0.466, *p* = 0.033) and in controls with a trend of significance (*R* = -0.384, *p* = 0.086), whereas the two correlations were not significantly different (*z*= -0.332, *p* = 0.740). There was a trend of significance that higher CAPS score correlated with lower left amygdala-vmPFC functional connectivity in PTSD participants (*R* = -0.404, *p* = 0.070) but not in PTSD participants (*R* = 0.262, *p* = 0.251), and the two correlations were significantly different (*z* = -2.311, *p* = 0.042). CAPS score was not related with brain activation in amygdala in either group (*p*-values > 0.1).

TLEQ score did not correlate with any brain response in either group (*p*-values > 0.1).

### 4. Behavioral and Brain Results (Two Groups without Age Difference)

The PTSD group was slightly older than the CONT group (*t*(48) = 2.143, *p* = 0.037) . To understand whether the between-group differences were confounded by age, we re-analyzed the data by excluding the oldest subject in the PTSD group as well as the youngest subject in the CONT group. The age of the remaining 24 subjects in the PTSD group (mean±SD = 42.9±9.6 years) were not significantly (*t*(46) = 1.702, *p* = 0.096) different from that of the 24 subjects in the CONT group (mean±SD = 38.7±7.6 years). As shown below, re-analyses in subjects with matched age showed comparable results relative to the findings reported in the main text, except that controls performed better (caught less) in nonthreat than threat context, and CAPS score was not significantly correlated with right amygdala-vmPFC functional connectivity in controls (*R* = -0.268, *n* = 24, *p* = 0.206).

#### 4.1 Task Performance

The average number of prey captures was non-significant for Group (*F*(1,46) = 2.130, *p* = 0.151), Context (*F*(1,46) = 0.464, *p* = 0.499), and Group x Context interaction (*F*(1,46) = 0.150, *p* = 0.700).

The average number of avatar captures was non-significant for Group (*F*(1,46) = 0.673, *p* = 0.416) and Context (*F*(1,46) = 1.676, *p* = 0.202), but significant for the Group x Context interaction (*F*(1,46) = 5.850, *p* = 0.020). The comparisons between group were non-significant for either threat (*t*(46) = -0.050, *p* = 0.961) or nonthreat (*t*(46) = 1.710, *p* = 0.094) contexts, whereas the between-context comparisons showed significance for the controls (*t*(23) = 2.418, *p* = 0.048) but not participants with PTSD (*t*(23) = -0.877, *p* = 0.778).

#### 4.2 Self-Report Questions

Threat-vs-nonthreat context elicited enhanced focus on avoiding predator (*F*(1,44) = 7.599, *p* = 0.008), increased initial (*F*(1,43) = 11.955, *p* = 0.001) and sustained anxiety(*F*(1,43) = 7.287, *p* = 0.010), and greater fear of being chased by the predator (*F*(1,43) = 6.928, *p* = 0.012).

Participants with PTSD compared to controls showed greater initial (*F*(1,43) = 6.028, *p* = 0.018) and sustained anxiety (*F*(1,43) = 9.538, *p* = 0.004) and a trend of significance of fear of being chased by predator (*F*(1,43) = 4.040, *p* = 0.051), but not for avoiding predator (*F*(1,44) = 0.001, *p* = 0.997).

There was no significant Group X Context interaction (*F-value*s < 1, *p-value*s > 0.3) for any self-report.

#### 4.3 Brain Activation in ROIs

Threat-vs-nonthreat context elicited larger activations across ROIs (*F*(1,46) = 4.850, *p* = 0.033). The main effect of ROI was also significant (*F*(2,92) = 9.549, *p* = 0.002). Further comparisons revealed that both left (*F*(1,46) = 11.627, *p* = 0.002) and right (*F*(1,46) = 9.071, *p* = 0.008) amygdala were related with larger brain activation than vmPFC, whereas there is no significant difference between left and right amygdala (*F*(1,46) = 0.435, *p* = 0.513). No result involving the effect of Group was significant (*F-value*s < 2.5, *p-value*s > 0.1).

#### 4.4 Amygdala-vmPFC Functional Connectivity

The significant Seed x Group interaction (*F*(1,46) = 5.005, *p* = 0.030) showed that PTSD patients versus controls were associated with smaller right amygdala-vmPFC functional connectivity than the left amygdala-vmPFC functional connectivity across contrast. However, the post hoc analyses did not find a significant between-group difference in either left (*F*(1,46) = 0.003, *p* = 0.960 uncorrected) or right (*F*(1,46) = 3.456, *p* = 0.069 uncorrected) amygdala-vmPFC functional connectivity. None of the other effects was significant (*F-value*s < 2.5, *p-value*s > 0.1).

#### 4.5 Brain-Performance Associations

For the threat-vs-nonthreat contrast, better performance (i.e. few avatar captures) was accompanied with larger vmPFC activation in PTSD participants (*R* = -0.544, *p* = 0.006) but not in controls (*R* = 0.228, *p* = 0.285), and the two correlations were significantly different (*z*= -2.728, *p* = 0.027). Better performance (i.e. few avatar captures) also related to stronger left-amygdala-vmPFC functional connectivity in controls (*R* = -0.411, *p* = 0.046) but not in PTSD participants (*R* = 0.227, *p* = 0.285), and the two correlations were different at trend level (*z*= -2.215, *p* = 0.053). The average number of avatar capture was related with neither brain activation in amygdala (*p*-values > 0.1) nor the left amygdala-vmPFC functional connectivity in either group (*p*-values > 0.1).

There was a trend of significance that the average number of prey capture was positively correlated with vmPFC activity (PTSD: *R* = -0.051, *p* = 0.813; CONT: *R* = 0.364, *p* = 0.080) and negatively correlated with left amygdala-vmPFC functional connectivity (PTSD: *R* = -0.008, *p* = 0.969; CONT: *R* = -0.384, *p* = 0.064) in controls but not in participants with PTSD. The correlations were not significantly different between-groups (*p*-values > 0.1). The average number of prey capture was related with neither brain activation in amygdala (*p*-values > 0.1) nor the right amygdala-vmPFC functional connectivity in either group (*p*-values > 0.1).

#### 4.6 Brain-Clinical Associations

For the threat-vs-nonthreat contrast, higher CAPS score was accompanied with smaller vmPFC activation in controls (*R* = -0.485, *p* = 0.024) but not in PTSD participants (*R* = 0.154, *p* = 0.471), and the two correlations were significantly different (Fisher’s *z*= -2.271, *p* = 0.046). Higher CAPS score was also associated with smaller right amygdala-vmPFC functional connectivity in PTSD participants (*R* = -0.482, *p* = 0.017) but not controls (*R* = -0.268, *p* = 0.206), whereas the two correlations were not significantly different (*p*-value > 0.1). CAPS score was related with neither brain activation in amygdala (*p*-values > 0.1) nor the left amygdala-vmPFC functional connectivity in either group (*p*-values > 0.1).

There was a trend of significance that higher TLEQ score was associated with lower right amygdala-vmPFC functional connectivity in controls (*R* = -0.361, *p* = 0.083) but not in participants with PTSD (*R* = 0.137, *p* = 0.524), and the two correlations were non-significantly different (*p*-value > 0.1). TLEQ score was related with neither brain activation in amygdala and vmPFC (*p*-values > 0.1) nor the left amygdala-vmPFC functional connectivity in either group (*p*-values > 0.1).

### 5. Brain Activation and Functional Connectivity in Inferior Frontal Gyrus (IFG)

Our previous study using the same task paradigm on healthy participants found that the threat > nonthreat contrast elicited stronger activation in bilateral IFG as well as greater right amygdala-bilateral IFG functional connectivity (12). Moreover, there was a trend for right amygdala-right IFG functional connectivity positively correlated with the average number of rewards earned during the threat condition only. Here, we also explored the between-group difference of activation and functional connectivity in bilateral IFG. The functionally defined ROIs of left (peak MNI coordinates -48,30,3) and right (peak MNI coordinates 48,43,-8) IFG were from our prior study using the same paradigm in non-clinical participants (12). As shown below, there is only one significant between-group difference, which is that the threat-vs- nonthreat contrast elicited larger activation in bilateral IFG in participants with PTSD compared to controls.

#### 5.1 Brain Activation

The mean beta values of BOLD activation in left and right IFG were entered into a Group (PTSD and CONT) by ROI (left IFG and right IFG) by Context (threat and nonthreat) repeated measures ANOVA model. There was a significant Group x Context interaction (*F*(1,48) = 4.732, *p* = 0.035), indicating that the threat-vs-nonthreat contrast elicited larger activation in bilateral IFG in participants with PTSD compared to controls. There was also a significant effect of ROI (*F*(1,48) = 5.589, *p* = 0.022), showing that left-IFG was associated with larger activation than right-IFG across contexts and groups. None of the other effects were significant (*F-value*s < 2.6, *p-value*s > 0.1).

#### 5.2 Amygdala-IFG Functional Connectivity

The mean beta values of left or right amygdala functional connectivity with either left or right IFG were entered into a Group (PTSD and CONT) by Seed (left amygdala and right amygdala) by Target (left IFG and right IFG) by Context (threat and nonthreat) repeated measures ANOVA model. There was a significant Group x Seed x Context x Target interaction (*F*(1,48) = 7.818, *p* = 0.007). Further analyses found significant Group x Seed x Context interaction by fixing the target area as the left IFG (*F*(1,48) = 17.499, *p* < 0.001) but not the right IFG (*F*-value < 1, *p*-value > 0.7). We further analyzed the Group x Context interaction for the functional connections with the left IFG, but did not find results surviving corrections (*p*-values > 0.1).

The main effect of Target also showed a trend of significance (*F*(1,48) = 3.349, *p* = 0.073) depicting that amygdala functional connectivity with left IFG is greater than that with right IFG.

None of the other effects was significant (*F-value*s < 2.3, *p-value*s > 0.1).

#### 5.3 Brain-Performance Associations

For the threat-vs-nonthreat contrast, there was a trend of significance that more avatar captures were associated with larger brain activity in the left IFG in PTSD (*R* = 0.346, *p* = 0.090) but not controls (*R* = 0.020, *p* = 0.925), and also a trend that more prey captures were associated with smaller right amygdala-left IFG functional connectivity in controls (*R* = -0.361, *p* = 0.076) but not in participants with PTSD (*R* = -0.025, *p* = 0.905).

No other correlation was found in either group between performance and brain responses (*p-*values > 0.1) to the threat-vs-nonthreat contrast.

#### 5.4 Associations of Brain-Clinical Measures

No significant correlation was found in either group between clinical measures (including CAPS score and TLEQ score) and brain responses (*p-*values > 0.2) to the threat-vs-nonthreat contrast.

### **Table 1S**. Psychotropic medications taken by PTSD patients daily.

| Patient ID | Medication 1 | Medication 2 | Medication 3 |
| --- | --- | --- | --- |
| 1 | Trazodone | Klonopin | Zoloft |
| 2 | Methylphenidate | Wellbutrin | - |
| 3 | Citalopram | Prozac | - |
| 4 | Clonazepam | Citalopram | - |
| 5 | Trazodone | - | - |
| 6 | Sertraline | - | - |
| 7 | Cymbalta | - | - |
| 8 | Venlafaxine | - | - |
| 9 | Mirtazapine | - | - |
| 10 | Sertraline | - | - |
| 11 | Citalopram | - | - |
| 12-25 | - | - | - |
